# Enhancing Carbon Conversion Efficiency and Product Yield Through Systematic Biocatalyst Design for Microbial Electrosynthesis

**DOI:** 10.64898/2026.09.22.753537

**Authors:** A. Abbas, M. Ahsanul Islam

**Affiliations:** Department of Chemical Engineering, Loughborough University, Loughborough, Leicestershire, LE11 3TU UK

**Keywords:** MES, commodity chemicals, CO_2_ reduction, bioelectrochemical systems, coulombic efficiency, and carbon conversion efficiency

## Abstract

The unprecedented rise in greenhouse gases such as carbon dioxide (CO_2_), and their detrimental effects on the atmosphere have intensified the current climate emergency. This challenge has inspired the development of novel carbon capture and utilisation (CCU) technologies, specifically microbial electrosynthesis (MES), for the bioelectrochemical fixation of CO_2_ into commodity chemical compounds (CCCs) using microbial biocatalysts. In this study, we systematically enriched and maintained robust mixed microbial communities of electroactive bacteria (EAB) from wastewater treatment sludge to serve as biocatalysts for MES. The MES performance of these biocatalysts under different operational conditions in bioelectrochemical systems (BES) reactors were investigated. BES reactors with enriched biocatalysts incubated and maintained under controlled conditions produced higher CCCs yields than those developed at ambient temperatures, while strict anaerobic operations further improved the MES efficiency compared to aerobic conditions. At an applied potential of −1000 mV vs Ag/AgCl, CO conversion reached up to 88.11%, significantly outperforming the lower potential (−600 mV) operation. These findings underscore the importance of systematic development of robust and stable biocatalysts, as well as maintaining strict anaerobic conditions, for achieving efficient MES performance, thereby demonstrating the potential of our approach for scalable industrial CCU applications.

## Introduction

The unprecedented rise in the emission of greenhouse gases such as carbon dioxide (CO_2_) from anthropogenic sources and their detrimental effects on the atmosphere have resulted in the current climate emergency^1^. Combating this climate emergency has inspired the development of a novel carbon capture and utilisation (CCU) technology, microbial electrosynthesis (MES) for bioelectrochemical fixation of CO_2_ into commodity chemical compounds (CCCs) using microorganisms as biocatalysts^2–4^.

MES is a developing field at the intersection of microbiology and electrochemistry, where microorganisms are employed to catalyse the conversion of electrical energy into chemical energy^5^. The MES process involves the use of electroactive bacteria (EAB) in a bioelectrochemical system (BES) reactor to perform extracellular electron transfer (EET) processes, during which EAB accept electrons directly or indirectly from an electrode (a cathode) and use these electrons to reduce CO into high-value CCCs such as organic acids^6,7^.

Although MES holds significant promise for sustainable chemical production and carbon sequestration, overcoming the technical, biological, and economic challenges is essential for its advancement from laboratory research to industrial-scale applications^8^. The cost of construction materials for BES reactors and peripherals, including working and counter electrodes, reference electrodes, and other relevant operational expenses must be reduced to make the MES technology economically viable and competitive with traditional chemical synthesis methods^9,10^.

Out of myriad operational challenges, higher electrical energy requirements are particularly crucial for the BES reactors to provide sufficient energy (i.e., reducing power) to anaerobic biocatalysts during MES and to maintain the optimum temperature for microbial growth. High energy requirements are also essential to remove oxygen (O_2_) from the reactor and its surrounding environment to ensure anaerobic biocatalysts’ growth during MES. Several studies reported various ranges of applied potentials for the operation of BES reactors^11–15^; these usually range from −500 to −800 mV vs standard hydrogen electrode (SHE) or −800 to − 1200 mV vs silver/silver chloride (Ag/AgCl) reference electrode. In addition, most of the previous studies have reported maintaining the reactor temperature between 30 and 70 °C, as this temperature range is essential for maintaining microbial health, activity, and productivity during the MES process. This temperature range is also desirable for maintaining enzyme activity, membrane fluidity, and accelerated growth rate of biocatalysts. However, the associated extra cost is inevitable, and these conditions can limit the operability of the BES reactors for MES applications^16–20^.

Another operational challenge is the requirement for longer reactor startup periods for the MES process to allow reactors and biocatalysts to reach at steady state^21^. This is crucial to guarantee the EAB interaction with electrodes, which is important to define MES performance and applicability^22^. In addition, coordination among EAB cells is advantageous to their growth, performance, and activity for CCCs production, and achieving this coordination requires a longer reactor startup time. Moreover, the lack of appropriate development, enrichment, and maintenance of biocatalysts further contributes to higher electrical energy requirements and a longer reactor startup time^23–25^. Since the EAB at the cathode is responsible for the reduction of CO_2_ into CCCs, the ability of EAB to reduce CO_2_ into CCCs in short intervals is paramount to reducing operational expenses and improving the MES efficiency^26–29^.

In this study, we aimed to address these challenges by investigating several parameters. Robust biocatalysts of EAB community were systematically developed, enriched and maintained from the activated sludge sourced from a wastewater treatment facility. MES experiments were then conducted under both aerobic and strictly anaerobic conditions across a range of applied potentials (−600 mV and −1000 mV vs Ag/AgCl). By evaluating the system performance under these varying conditions, this study provides insight into the durability and effectiveness of BES for CO conversion into CCCs during MES. The findings highlight the critical role of optimised EAB and BES operating conditions in achieving a robust and efficient MES performance, underscoring their potential for scalable industrial CCU applications.

## Materials and Methods

### Chemicals and media preparation

All chemicals were purchased from Fisher Scientific (Loughborough, UK) unless otherwise specified. The growth medium, also used as catholyte and anolyte, was prepared based on Yang et al^30^ with the following modifications: the base medium (per litre) contained 0.35 g KH_2_PO_4_, 0.25 g K_2_HPO_4_, 0.50 g KCl, 0.25 g NH_4_Cl, 0.60 g MgCl_2_·6H_2_O, 0.16 g CaCl_2_·2H_2_O, and 1.20 g NaCl and was supplemented with trace elements (1 mLL ¹), sodium 2-bromoethanesulfonate (5 mM), tungstate–selenium solution (0.1 mL L ¹), vitamin solution (2.5 mL L ¹), and nutrient broth (13 g L ¹; peptone and yeast extract) for heterotrophic growth. For autotrophic conditions, the nutrient broth was omitted and replaced with CO_2_. All media were sterilized at 121 °C for 20 minutes, followed by purging with N_2_ for 30 minutes to establish anaerobic conditions. The pH was adjusted to 7.0 using 1 M HCl or NaOH.

### Biocatalyst development, enrichment, and maintenance

Biocatalysts were developed and enriched by adapting procedures described in Logrono et al^31^ and shown in Figure 1. First, 500 mL of distilled water in a 1 L flask was boiled and cooled below 30 °C while continuously flushed with N_2_. Next, 100 g of sludge was added to the flask to create a mixed microbial culture, followed by exchanging the flask’s headspace with N_2_ and then closed with a butyl rubber septum. In parallel, four 250 mL serum bottles were prepared anaerobically, filled with 50 mL heterotrophic medium, and sealed with butyl rubber septa. The bottles were placed in a 100 °C water bath for 20-30 minutes, followed by purging the headspace with an anaerobic gas mixture (80% N_2_ and 20% CO_2_; BOC Gases, UK)^32^.

**Figure 1.**
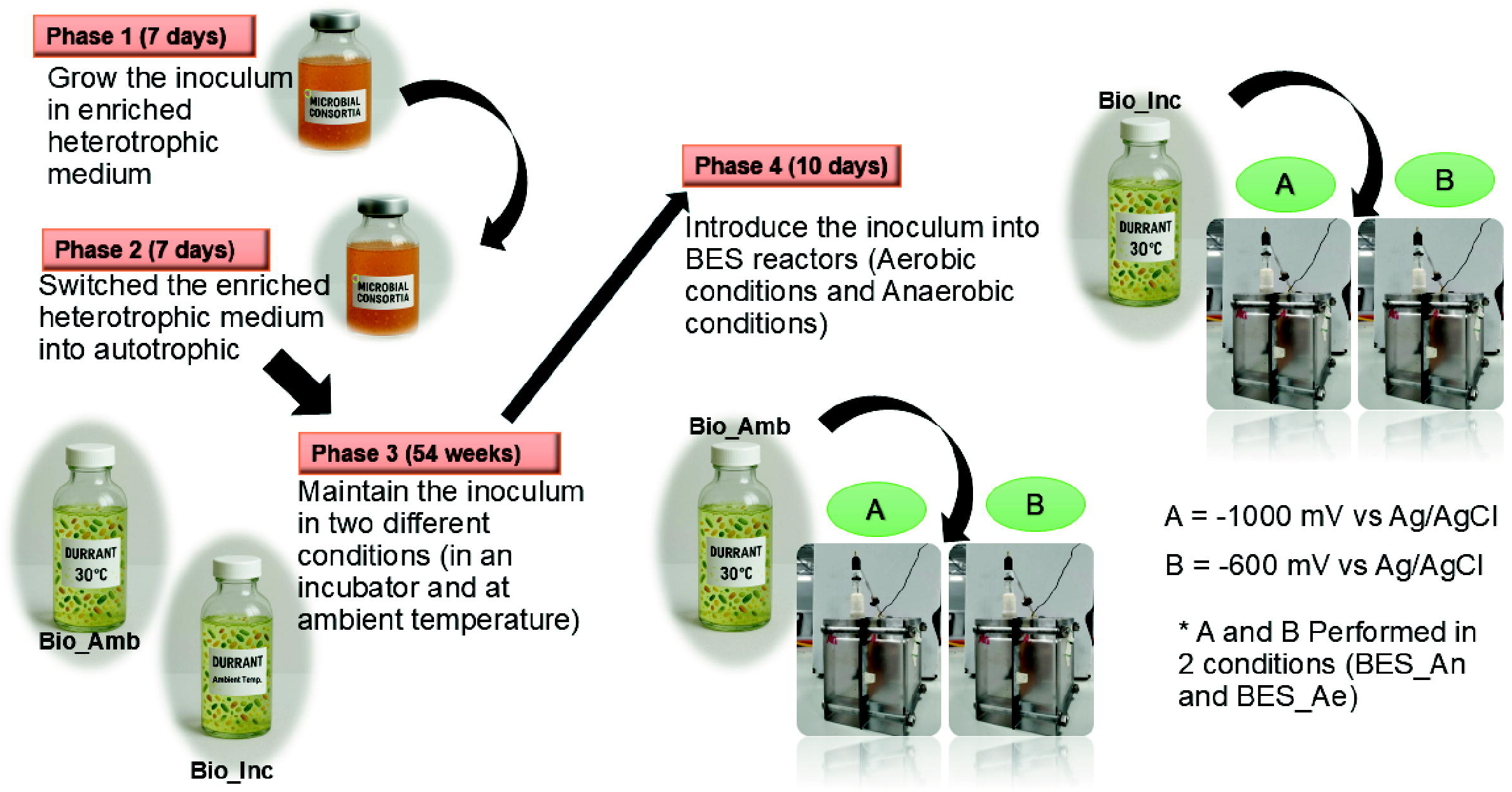
Flow chart of enrichment, maintenance, and microbial electrosynthesis experiments. Maintenance was performed at two different temperatures: constant 30 °C (incubator) and ambient temperatures (18–25 °C). Microbial electrosynthesis experiments were performed under two different conditions (aerobic and anaerobic).

Three bottles were inoculated with cultures from the flask, where a 10 mL inoculum was used for each of the three bottles while the fourth bottle was used as a control (without any cultures in it). All bottles were placed inside a vinyl anaerobic chamber (Coy Laboratory Products, Inc., Michigan, US) to maintain strict anaerobic conditions. The bottles were incubated at 30 °C in a shaker incubator at 120 rpm for 30 minutes. The cultures were grown heterotrophically for 7 days (Phase 1 in Figure 1), followed by inoculating three 250 mL serum bottles containing the autotrophic growth medium and using a 10 mL inoculum from the Phase 1 cultures (Phase 2 in Figure 1). After 7 days, these autotrophic cultures were used to create two sets of biocatalysts: Bio_Amb and Bio_Inc, where Bio_Amb cultures were maintained at ambient temperatures (20-30 °C) and Bio_Inc were maintained in an incubator at 30 °C. Both cultures were maintained inside the vinyl anaerobic chamber by bubbling 263.9 mg. L^-1^ of CO_2_ for 30 minutes^33^ every week for 54 weeks (> 1 year) (Phase 3 in Figure 1). These Phase 3 cultures were then used in BES reactor experiments (Phase 4 in Figure 1).

### BES reactor construction and operation

BES reactor components were obtained from Fuel Cell Store (TX, USA). Reactors were constructed from Perspex with external dimensions of 8.5 × 9.0 × 3.5 cm (W × H × D). The anodic and cathodic chambers (5.5 × 5.5 × 3.0 cm each) were separated by a proton exchange membrane (Fumasep FKB CEM PEEK exchange membrane; thickness: 110 – 140 μm (dry); weight per unit area: 10 – 13 mg cm^-2^; ion exchange capacity: 0.8 – 1.0 meq g^-1^). Membranes were pretreated in 0.5 M NaCl at 25 °C for 24 hours.

A PTFE-treated (10%) carbon cloth gas diffusion layer (4 × 5 cm^2^; thickness 0.41 mm; 200 g m^-2^) was used as cathode while a titanium mesh (3 × 4 cm²) was used as anode. Electrodes were connected using titanium wires. The Ag/AgCl reference electrode was calibrated at +0.209 V versus the SHE. Unless otherwise specified, all applied potentials mentioned in this study are measured against the Ag/AgCl reference electrode^2^.

Salt bridges were prepared using 2.0 g agar and 8.7 g NaCl dissolved in 50 mL deionised water. The molten gel (3 mL) was transferred into pipette tips, solidified, and sealed after inserting the reference electrode. Electrochemical measurements were conducted using a four-channel potentiostat (Whistonbrook, UK) in a three-electrode configuration. Chronoamperometry was applied to maintain constant potentials, and current was continuously recorded.

BES reactors were operated under aerobic (BES_Ae) and strictly anaerobic (BES_An) conditions. Anaerobic experiments were performed inside the vinyl anaerobic chamber. All experiments were conducted in triplicate using −1000 mV and −600 mV (vs Ag/AgCl) applied potentials. The electrolyte composition is the same as the growth medium, except containing nutrient broth and yeast extract. The catholyte was sparged with CO (30 min; 263.9 mg L ¹), and the anolyte was sparged with N (30 min). Control experiments included (i) an abiotic reactor operated at −1000 mV and (ii) biotic reactors containing Bio_Amb and Bio_Inc cultures and operated under open-circuit potential (OCP).

### Sequencing and bioinformatic analysis

DNA was extracted from the samples using the DNeasy PowerWater kit (14900-100-NF, QIAGEN, Germany) following the manufacturer’s protocol. The extracted DNA samples were sequenced at EnviSion BioSequencing and BioComputing facility at the University of Birmingham using the Illumina MiSeq platform, and the raw data was deposited at NCBI BioProject database (Accession number, PRJNA1427088). The raw sequence data (fastQ files) was analysed using the following workflow: raw paired-end sequencing reads were processed using the DADA2 pipeline (v1.28.0) in R following the standard workflow described by Callahan et al^34^; quality filtering, dereplication, denoising, merging of paired-end reads, and chimera removal were performed to infer high-resolution amplicon sequence variants (ASVs). Taxonomic assignment was carried out using the ‘assignTaxonomy()’ function with the Silva v138 reference database^35^. The resulting ASV count tables and taxonomic classifications were exported as CSV files for further downstream analysis.

### Sampling and analyses

Temperature in the anaerobic chamber and BES reactors was monitored using a traceable digital thermometer (Fisher Scientific, Loughborough UK), and pH was measured with Hanna digital instruments (Fisher Scientific, Loughborough UK). OD_600_ was measured using a UV–vis spectrophotometer (UV-mini-1240, Shimadzu, Japan) on 4 mL liquid samples withdrawn via syringe and cannula. Volatile fatty acids (formate, acetate, propionate, iso-butyrate, butyrate, isovalerate, valerate, and hexanoate) in catholytes were quantified by ion chromatography (Eco IC, Metrohm, Switzerland) equipped with a Metrohm 6.1005.200 column and autosampler. Samples were filtered, diluted 1:1 with deionised water, acidified with 20 μL of concentrated HCl (pH < 2), and analysed using seven-point calibration standards.

Alcohols (methanol, ethanol, isopropanol, butanol, and hexanol) and headspace gases were analysed using gas chromatography (GC-2010 Tracera, Shimadzu, Japan) equipped with a BID detector, autosampler, Zebron ZB-WAXplus capillary column (Phenomenex), Shin Carbon ST micro packed column (80/100, Restek), and helium as the carrier gas. Filtered liquid samples (1.5 mL) were sealed in vials for analysis, and 15 mL of headspace gas was injected into the inlet. Statistical differences were evaluated using a one-tailed t-test.

### Calculation method

The acetate production rate (R_acetate_) was calculated using the following equation and adopted from Faraghiparapari and Zengler^36^:

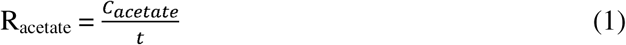

where C_acetate_ (mg. L^−1^) and t (d) are the acetate concentration and time, respectively.

The carbon dioxide conversion efficiency (CO_2_CE) was calculated according to Li et al^37^ using the following equation:

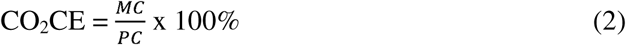

where PC is the carbon concentration (mg. L^−1^) in the electrolyte (in the form of CO_2_) and MC (mg. L^−1^) is the carbon concentration in the CCCs (mg. L^−1^), respectively. The Coulombic efficiency (CE) was calculated as the moles of electrons recovered into CCCs using the method adapted from Li et al^37^:

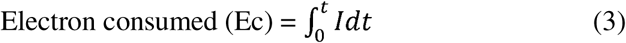

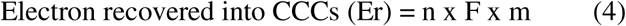

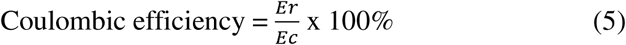

where I is the electron consumption recorded by the potentiostat, t is the operation time of the BES reactor, F is the Faraday’s constant (96485 C/mol of e^-^), n is the number of moles of electrons recovered into CCCs, and m is the number of moles of CCCs produced during the experiment.

## Results and Discussion

### Development and characterisation of stable electroactive biocatalysts

Two electroactive biocatalysts (Figure 1), designated as Bio_Amb (maintained at ambient temperatures, 18–25 °C) and Bio_Inc (maintained at constant 30 °C in an incubator), were systematically enriched from activated sludge through a multi-stage protocol (see materials and methods) designed to select for EAB capable of extracellular electron uptake and autotrophic CO_2_ fixation. Before deployment in BES reactors, mixed cultures containing EAB were cultivated heterotrophically and autotrophically through three consecutive phases in nutrient-rich and nutrient-free medium, respectively (Figure 1). The medium was supplemented with 2-bromoethanesulfonate to selectively suppress methanogenesis, thereby preventing competition for electrons and diversion of carbon to methane formation. Microbial growth was monitored via measuring OD_600_ (Figure 2) for heterotrophic (Figure 2A) and autotrophic (Figure 2B) conditions, and methane suppression was confirmed by monitoring headspace gas composition using gas chromatography (data not shown).

**Figure 2.**
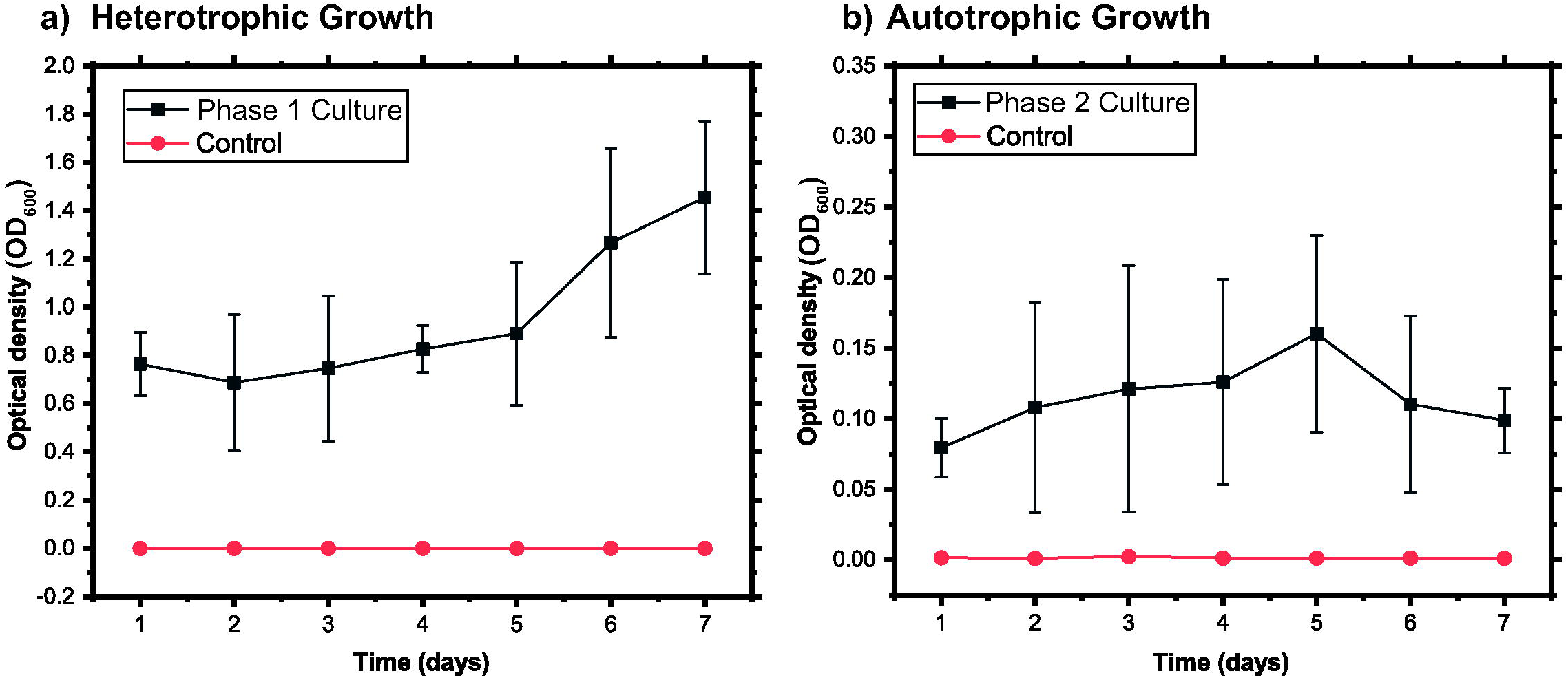
Optical density at 600 nm (OD) of biocatalysts grown under different culture conditions: (a) grown in heterotrophic medium and (b) grown in autotrophic medium. Error bars represent standard deviation (SD) of triplicate experiments (n = 3).

To confirm the metabolic activity of biocatalysts before their deployment in BES reactors, organic acid production by the mixed cultures was quantified during the maintenance phase by ion chromatography. Both Bio_Amb and Bio_Inc predominantly produced formate and iso-butyrate (Table S1 in the Supplementary Information), while the formate production rate (13.3 ± 4.1 mg L^-1^ week^-1^) was faster for Bio_Inc than Bio_Amb but the iso-butyrate production rate showed the opposite trend (9.1 ± 3.7 mg L^-1^ week^-1^). This difference in product formation rates can be attributed to the differences in microbial community structures originated from maintaining the cultures at different conditions and corroborated through metagenome sequencing analyses described later. Sustained organic acid formation and detection across both biocatalysts confirmed the establishment of metabolically active, acetogenic mixed microbial communities capable of fixing CO_2_ into organic acids using available reductants (i.e., H_2_). The product profiles are consistent with prior reports of mixed acetogenic cultures, including *Sporomusa*, *Desulfovibrio*, and *Clostridium* producing short and medium-chain carboxylates when supplied with adequate reducing equivalents^5, 30, 38, 39^ with the production of longer-chain products was particularly attributed to chain-elongating *Clostridium* species^40^.

The two-stage enrichment strategy employed in this study by combining heterotrophic and autotrophic pre-growth with methanogen suppression, followed by temperature-differentiated maintenance inside a vinyl anaerobic chamber (Figure S1 in the Supplementary Information), constitutes a systematic biocatalyst design framework. Rather than relying on undefined environmental inoculation usually employed for MES applications^41,42^, this protocol specifically helps modulating the community composition and metabolic bias before deploying the cultures in BES reactors for MES purposes. The resulting mixed communities of EAB exhibited markedly different microbial community structures and product profile performance, as described in detail in the following sections, in addition to providing an experimental platform to decouple and analyse the effects of community composition from operational parameters in BES reactors during the MES process.

### Organic acid production performance of biocatalysts under aerobic and strictly anaerobic conditions in BES reactors

#### Aerobic BES reactors

Under aerobic conditions, BES reactors (BES_Ae) equipped with Bio_Amb and Bio_Inc biocatalysts at two different applied potentials exhibited a characteristic temporal shift in the dominant organic acid production profiles (Figure 3). Iso-butyrate was the primary product on day 1 across all BES_Ae configurations at both −1000 mV and −600 mV (vs. Ag/AgCl) applied potentials, while acetate was detectable only at trace levels. Over the 10-day experimental period, iso-butyrate concentration declined progressively while acetate was accumulated, ultimately becoming the dominant product (Figures 3A, 3B, 3C, and 3D). Although different biocatalysts were employed in BES reactors operating in aerobic conditions, no statistically significant difference (t-test, p > 0.05) in product profiles was observed between BES_Ae+Bio_Amb (Figures 3A and 3B) and BES_Ae+Bio_Inc (Figures 3C and 3D) configurations at either potential.

**Figure 3.**
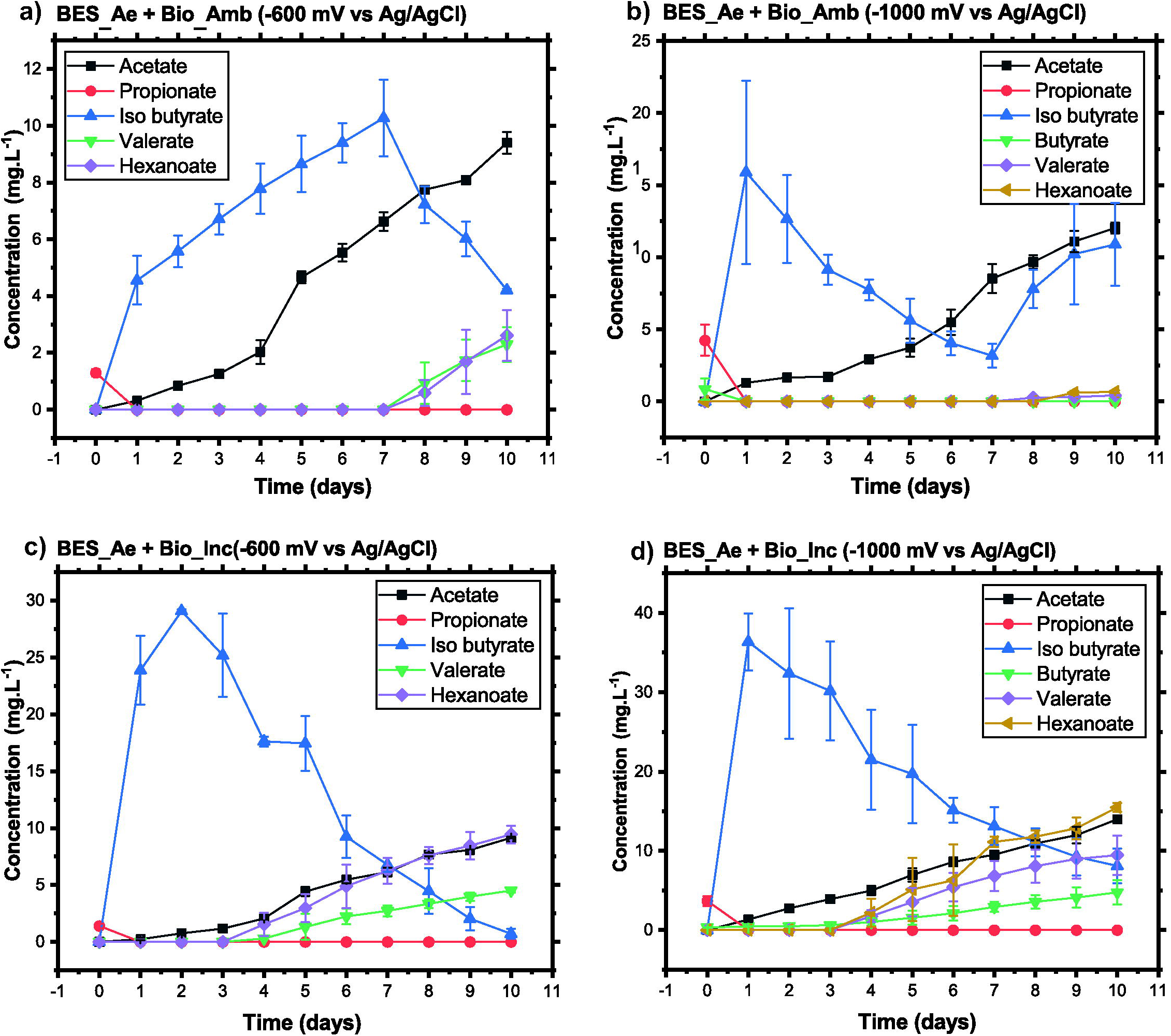
Product profiles of MES experiments in BES under aerobic conditions. (a) BES_Ae + Bio_Amb at −600 mV vs. Ag/AgCl; (b) BES_Ae + Bio_Amb at −1000 mV vs. Ag/AgCl; (c) BES_Ae + Bio_Inc at −600 mV vs. Ag/AgCl; and (d) BES_Ae + Bio_Inc at −1000 mV vs. Ag/AgCl. Samples were collected at the same time each day. Error bars represent the standard deviation (SD) of triplicate experiments (n = 3).

The early-phase dominance of iso-butyrate production can be mechanistically attributable to transient microaerophilic conditions arising from oxygen intrusion during the reactor setup and O_2_ evolution in the anode^43^. Under these conditions, obligate anaerobes such as *Clostridium spp*. are transiently suppressed, permitting facultative or oxygen-tolerant bacteria, including *Bacillus*, *Pseudomonas*, and *Acinetobacter*, to ferment branched-chain amino acids and generate iso-butyrate ^43^.

As residual O_2_ was progressively consumed by facultative and oxygen-tolerant populations (e.g., *Bacillus, Pseudomonas, Acinetobacter*) during the initial days, the reactor environment transitioned toward anaerobiosis, favouring the establishment of obligate anaerobic homoacetogens such as *Clostridium spp*. and *S. ovata*-type biocatalysts. These organisms possess electron-bifurcating hydrogenases that oxidise cathodically generated H_2_ to yield reduced ferredoxin and NADH, the electron carriers driving CO_2_ fixation via the WL pathway^44^. This microbial succession, from fermentative aerobes/ facultative anaerobes utilising branched-chain amino acids to H_2_/ CO_2_-utilising homoacetogens, accounts for the progressive decline in iso-butyrate and the concomitant, sustained rise in acetate over the 10-day experiment period.

Consistent with this proposed mechanism, acetate accumulation was more pronounced at the more negative applied potential (12.04 mg/L at −1000 mV vs. 9.40 mg/L at −600 mV by day 10, corresponding to average production rates of 1.20 and 0.94 mg L^-1^ day^-1^, respectively, a ∼28% enhancement at the higher cathodic driving force). This trend supports the interpretation that increased H_2_ evolution at more negative potentials accelerated the depletion of residual O_2_ and promoted earlier colonisation by H_2_-oxidising homoacetogens, consistent with the potential-dependence of cathodic hydrogen evolution described for O_2_ intrusion in BES systems ^43^. Notably, the rate of acetate accumulation slowed toward the end of the experimental period at both potentials, despite the absence of any decline in concentration. This deceleration is consistent with a shift toward a substrate (H_2_/CO_2_)- limited, quasi-steady-state regime as biomass increased relative to electron supply^38^, a pattern also linked to feedback inhibition of acetogenesis at elevated acetate concentrations in related anaerobic systems^45^, rather than a shift toward a competing metabolic pathway such as chain elongation.

Despite this positive trend, the acetate production rates observed in BES_Ae reactors (R_acetate = 0.94–1.20 mg L^-1^ day^-1^) remained substantially lower than those typically reported for optimised *S. ovata*-based MES systems using carbon cloth cathodes under continuous, fully anaerobic poised-potential operation, that range from approximately 0.1 to 0.7 g L^-1^ day^-1^ ^46^. This gap of roughly two to three orders of magnitude is consistent with the transient aerobic start-up conditions specific to BES_Ae reactors, which likely impose an initial performance penalty relative to systems inoculated and operated entirely under anaerobic conditions from the outset, rather than reflecting an intrinsic limitation of the biocatalysts themselves^47^.

#### Strictly anaerobic BES reactors

BES reactors with Bio_Inc and Bio_Amb in strictly anaerobic conditions (BES_An) were operated under −1000 mV and −600 mV applied potentials inside a vinyl anaerobic chamber, eliminating the oxygen-mediated interferences observed previously under aerobic conditions. Two BES reactors operated under OCP and same anaerobic conditions with biocatalysts produced trace rate of acetate and propionate (Table S2 in the Supplementary Information). These results are attributable to H_2_-driven CO_2_ reduction by the biocatalysts in the absence of electrochemical driving force, confirming their inherent acetogenic activity^48, 49^. No products were detected in abiotic controls (Table S2 in the Supplementary Information), confirming that organic acid formation was biologically mediated.

Under the applied potentials, acetate was the dominant product in all BES_An reactors, as shown in the product profiles (Figure 4). BES_An+Bio_Amb produced acetate at 177.03 ± 30.24 mg L ¹ (R_acetate = 17.70 mg L^-1^ d^-1^) at −1000 mV and 90.1 ± 11.74 mg L^-1^ (R_acetate = 9.75 mg L^-1^ d^-1^) at −600 mV (Figures 4A and 4B). BES_An+Bio_Inc achieved markedly superior performance (Figures 4C and 4D), with acetate reaching 394.21 ± 4.01 mg L^-1^ (R_acetate = 39.20 mg L^-1^ d^-1^) at −1000 mV and 269.21 ± 4.66 mg L^-1^ (R_acetate = 26.42 mg L^-1^ d^-1^) at −600 mV. The approximately 2.2-fold higher acetate output of Bio_Inc relative to Bio_Amb under identical electrochemical conditions is consistent with the enrichment of *Sporomusa ovata* in the Bio_Inc community, a highly electrotrophic acetogen recognised as the most productive pure-culture MES biocatalyst, capable of producing 51.1 g m^-2^ d^-1^ of acetate^10, 50^. This observation was also confirmed by metagenome sequencing analysis discussed later. Notably, the acetate production rate achieved by BES_An+Bio_Inc (39.20 mg L ¹ d ¹ at −1000 mV) compares favourably with those reported for mixed-culture MES systems operating with similar carbon-based cathodes such as the 49.2 mg L ¹ d ¹ reported by Mateos et al^51^, despite the use of unmodified carbon cloth and without implementing headspace gas recirculation or any other advanced reactor configurations in BES_An+Bio_Inc. BES_An+Bio_Inc at −1000 mV also exhibited the greatest product diversity, with up to seven organic acids detected (Figure 4D), reflecting active chain-elongation pathways in operation under high cathodic reducing power. This diversity in product formation was narrowed to four compounds at − 600 mV (Figure 4C), suggesting that higher applied potentials support a more energetically diverse metabolic network, likely through generating enhanced H_2_ partial pressure at the cathode surface. BES_An+Bio_Amb showed a more constrained but stable product range (four to five compounds) at both applied potentials (Figures 4A and 4B). The differences in product formation performance observed between BES reactors with both biocatalysts operated under aerobic and anaerobic conditions are statistically significant (p < 0.01; Table S3 in the Supplementary Information), suggesting that strictly anaerobic conditions and high reducing power supply significantly enhanced the MES performance by developed biocatalysts.

**Figure 4.**
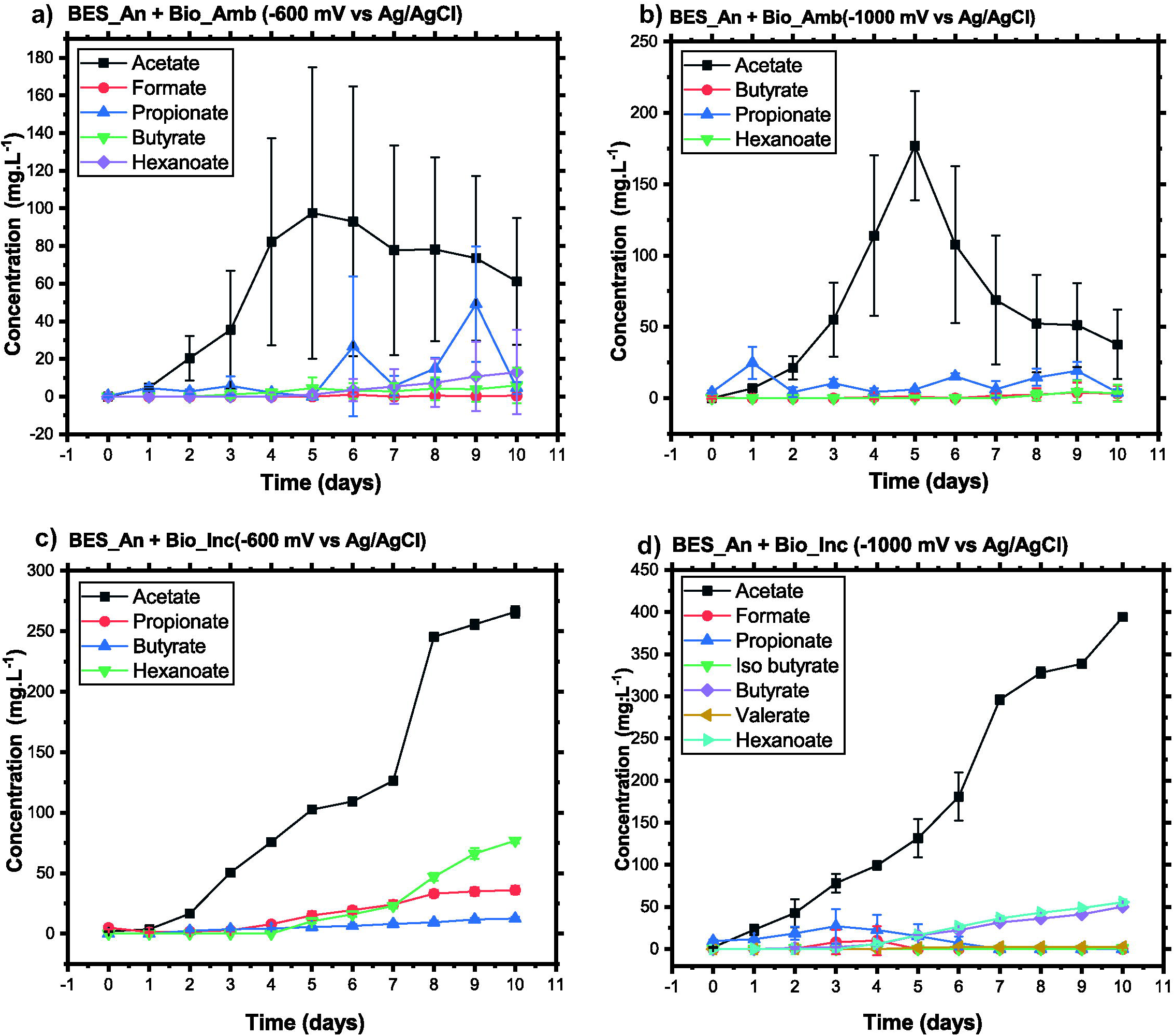
Product profiles of MES experiments in BES under strictly anaerobic conditions. (a) BES_An + Bio_Amb at −600 mV vs. Ag/AgCl; (b) BES_An + Bio_Amb at −1000 mV vs. Ag/AgCl; (c) BES_An + Bio_Inc at −600 mV vs. Ag/AgCl; and (d) BES_An + Bio_Inc at −1000 mV vs. Ag/AgCl. Samples were collected at the same time each day. Error bars represent the standard deviation (SD) of triplicate experiments (n = 3).

### Electron transfer mechanisms and their implications for MES performance

Although direct electrochemical characterisation (e.g., cyclic voltammetry, electrochemical impedance spectroscopy) was not performed in this study, the performance data presented here, in conjunction with community composition analysis (discussed later), allow mechanistic inferences about the dominant electron-transfer pathways operative in each reactor configuration. In MES systems, electron uptake by microorganisms generally proceeds via three non-mutually exclusive mechanisms: (i) direct electron transfer (DET)^52, 53^, in which bacteria physically contact the electrode and accept electrons via outer membrane cytochromes or conductive biofilm networks; (ii) mediated electron transfer (MET)^54, 55^, in which soluble mediators shuttle electrons between the electrode and planktonic cells; and (iii) H_2_-mediated indirect electron transfer^56^, in which cathodically evolved H_2_ acts as a soluble electron carrier oxidised by hydrogenase enzymes at the microbial cell surface.

The dominance of H_2_-mediated transfer is strongly supported by the experimental conditions used in this study. The anaerobic chamber atmosphere contained 2.5–4% H_2_, and the applied cathodic potentials (−600 to −1000 mV vs. Ag/AgCl) are thermodynamically favourable for H_2_ evolution on carbon surfaces. The detection of acetate and propionate in OCP reactors exposed to the H_2_-containing chamber atmosphere confirms that both biocatalysts can actively consume H_2_ for CO_2_ reduction in the absence of any applied potential (Table S2 in the Supplementary Information). This is consistent with the well-established hydrogenase activity of *S. ovata* and *Clostridium spp*., which oxidise H_2_ to generate reduced ferredoxin and NADH, the key electron carriers for the Wood–Ljungdahl (WL) pathway^57^.

The WL pathway is the core metabolic engine for acetate production in both biocatalysts. In this pathway, two molecules of CO are sequentially reduced: one to formate (via formate dehydrogenase) and subsequently to CO (via CO dehydrogenase), and the other to a methyl group (via the methyl branch), with both products converging at the acetyl-CoA synthase complex to yield acetyl-CoA, which is then phosphorylated to acetate with coupled ATP generation^58, 59^. The transient accumulation of formate detected in both Bio_Amb and Bio_Inc during the maintenance phase (Table S1 in the Supplementary Information) is consistent with this intermediate being a natural WL pathway product and a secondary electron sink during suboptimal growth conditions^60, 61^.

The superior CE and CO CE achieved by Bio_Inc under anaerobic conditions are mechanistically consistent with the presence of *S. ovata* (9.45% relative abundance), a bacterium with high affinity for both cathodic electrons and H_2_ as electron donors, coupled with a highly efficient WL pathway. *S. ovata* possesses a novel *Sporomusa*-type electron-bifurcating transhydrogenase (Stn) that links NADH and ferredoxin pools, enabling efficient NADP reduction, a bottleneck in many acetogens^26^. This enzymatic capacity likely contributes to the higher electron recovery and reduced side-product formation observed in BES_An+Bio_Inc. The progressive enrichment of *S. ovata* and *Clostridium spp*. on the cathode surface under strict anaerobiosis (discussed later) further suggests that biofilm-mediated DET may contribute alongside H_2_-mediated transfer, though this requires electrochemical confirmation in future studies^62–65^.

### CO conversion efficiency, coulombic efficiency, and contextual benchmarking

Table S4 in the Supplementary Information summarises the key performance metrics of the BES reactors from this study alongside relevant literature data. CO_2_CE and CE were evaluated as primary indicators of system efficiency, reflecting inorganic carbon utilisation and electron recovery into target products, respectively. The calculated values of CO_2_CE and CE from the present experiments are shown in Figure 5, and the recorded data are presented in Tables S5 –S12 in the Supplementary Information.

**Figure 5.**
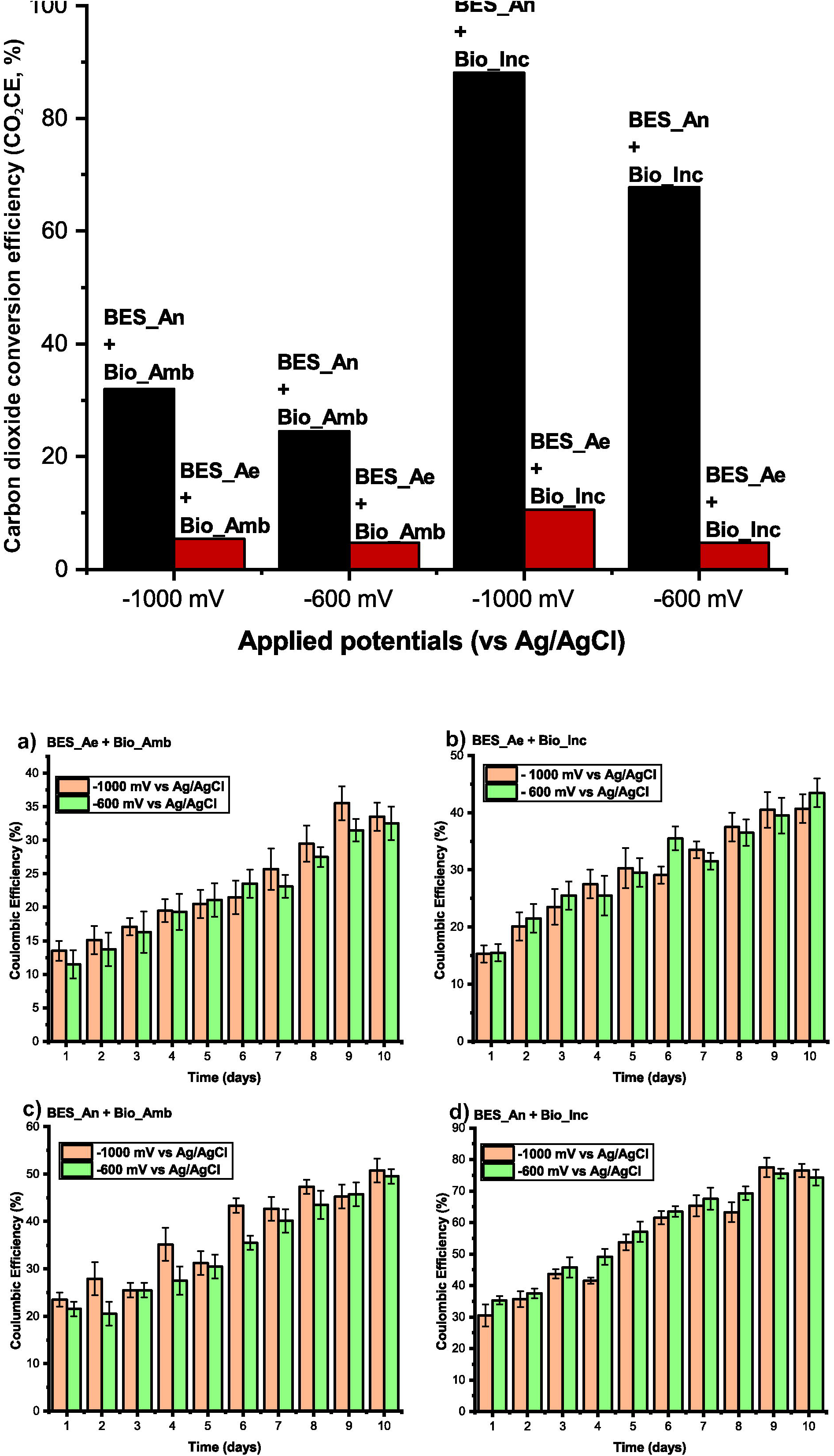
(Top) Carbon dioxide conversion efficiency (CO_2_CE) in BES reactors with different biocatalysts at −1000 and −600 mV vs. Ag/AgCl. (Bottom) Coulombic efficiency (CE) under the same conditions: (a) BES_Ae+Bio_Amb, (b) BES_Ae+Bio_Inc, (c) BES_An+Bio_Amb, and (d) BES_An+Bio_Inc. Error bars represent standard deviation (SD) of triplicate experiments (n = 3).

Under aerobic conditions, CO_2_CE was substantially limited, with BES_Ae+Bio_Amb and BES_Ae+Bio_Inc achieving maximum conversions of 5.40% and 10.55% at −1000 mV, respectively (Figure 5A). This limited carbon conversion efficiency reflects the well-established inhibitory effects of O_2_ on the WL pathway in obligate acetogens, cathodic competition between CO_2_ and O_2_ as electron acceptors, and community-level shifts away from obligate anaerobes^43, 47, 60, 66^.

In contrast, strictly anaerobic operations yielded a markedly superior performance. BES_An+Bio_Amb achieved CO_2_CE of 31.98% (−1000 mV) and 24.50% (−600 mV), while BES_An+Bio_Inc attained 88.11% (−1000 mV) and 67.78% (−600 mV), representing near-complete utilisation of dissolved inorganic carbon at the higher applied potential (Figure 5A). CE followed a consistent pattern: aerobic reactors achieved 34–42% (Figures 5B and 5C) while anaerobic reactors reached 49–51% (Figure 5D) and 79–80% (Figure 5E) coulombic efficiency. These values exceed those reported by Tahir et al. (41% CE)^67^ and Zhang et al. (76% CE)^46^, both of which employed the same untreated carbon cloth cathode, confirming that the performance gains achieved in this study are attributable to biocatalyst design and operational environment rather than using modified electrodes.

Contextualising these results within the broader MES literature (Table S4) reveals that while advanced reactor designs such as NanoWeb-RVC electrodes^56^, flow-electrode systems^68^, and H_2_-mediated high-titre reactors^69^ can achieve substantially higher acetate production rates (up to 12.5 g L^-1^ in 14 days) through electrode modification or reactor engineering^70^, the CE achieved by BES_An+Bio_Inc (80%) is competitive with many of these systems and was obtained using unmodified carbon cloth, a considerably simpler and more cost-accessible approach. Furthermore, the acetate production rates reported here are comparable to those of mixed-culture systems employing more complex reactor configurations, such as the continuous headspace recirculation system of Mateos et al^71^, reinforcing the contribution of biocatalyst design as a primary performance determinant for MES. This outcome positions community-level biocatalyst engineering as an underexplored but highly effective lever for improving MES performance without capital-intensive electrode modification efforts.

Cathodic current consumption was consistently higher (represented by higher CE) at −1000 mV than at −600 mV, confirming that more negative potentials enhance electron uptake rates. Despite similar current densities between BES_An+Bio_Inc and BES_An+Bio_Amb (p > 0.05), the former generated substantially higher CCC titres (Figures 4C and 4D), a discrepancy attributable to a superior CE rather than a greater current draw. This dissociation between current density and product yield underscores the importance of microbial electron utilisation efficiency as a determinant of BES productivity. The feasibility of operating at ambient temperatures (Bio_Amb) while still achieving CE of 51% and CO_2_CE of 31.98% further demonstrates that strict anaerobiosis, rather than elevated temperature, is the dominant driver of system efficiency, with important implications for energy-affordable scale-up of MES systems.

### Microbial community composition and mechanistic links to BES performance

Metagenomic sequencing of Bio_Amb and Bio_Inc biocatalysts, as well as BES reactor communities in aerobic and anaerobic conditions (Figure 6) provided a deeper mechanistic insight into the performance differences observed across experimental conditions. Bio_Amb was dominated by *Clostridium sensu* stricto 12 (16.67%), *Stenotrophomonas* (16.55%), *Clostridium* (15.38%), *Oscillospira* (12.57%), and *Clostridium sensu* stricto 4 (11.39%). Bio_Inc exhibited a distinct community architecture dominated by *Clostridium sensu* stricto 10 (10.37%), *S. ovata* (9.45%), *Clostridium sensu* stricto 12 (8.73%), and *Clostridium* (7.23%). These compositional differences directly reflect the influence of maintenance temperature on microbial community selection: a constant 30°C incubator temperature preserved mesophilic acetogens including *S. ovata*, whereas maintenance at ambient temperatures (18–25°C) favoured psychrotolerant and stress-adapted taxa such as *Psychrobacter*, *Pseudomonas*, and *Bacillus*, microorganisms with lower electrotrophic potential^72^.

**Figure 6.**
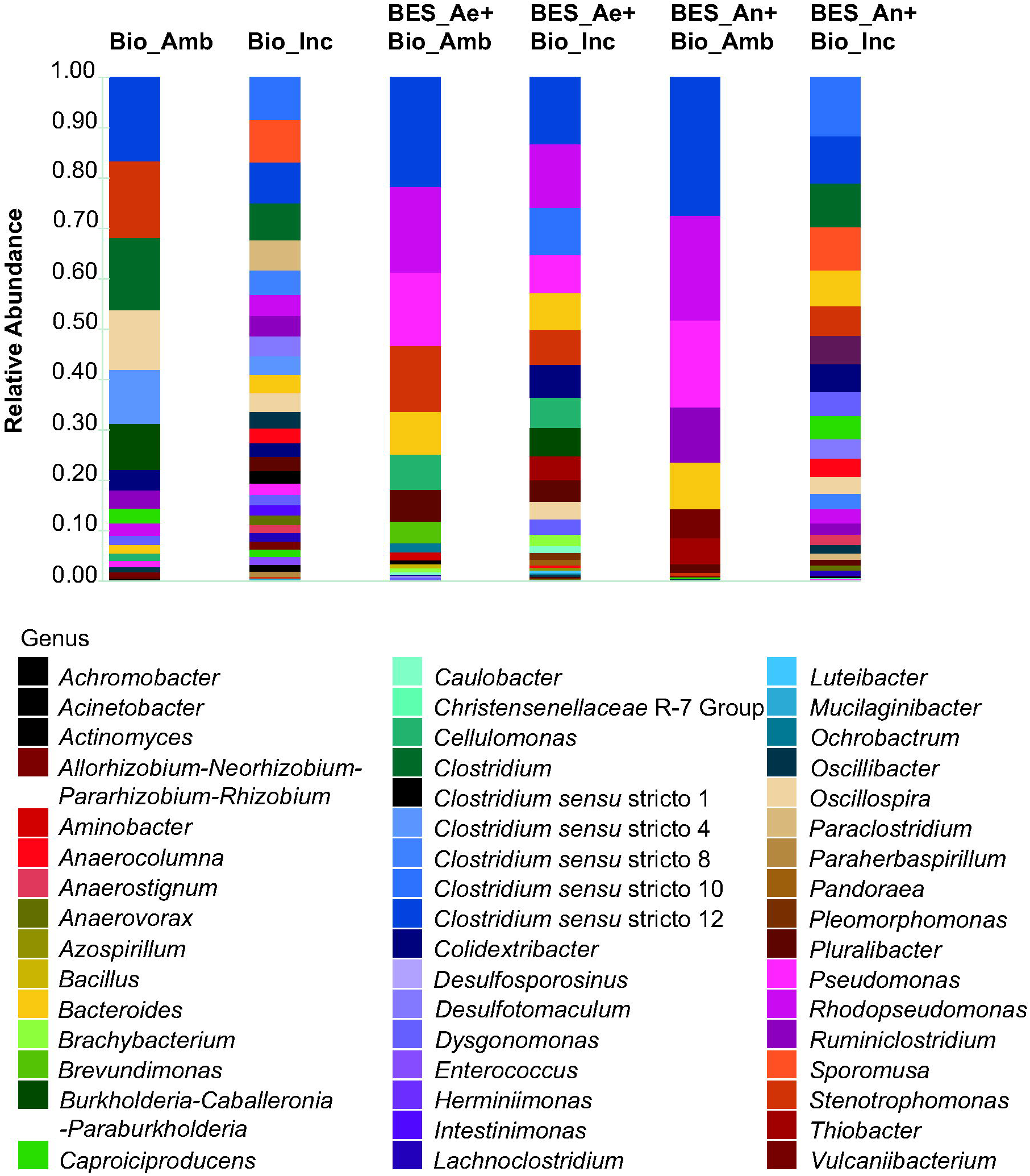
Relative abundance of microbial communities at the genus level from biocatalysts maintenance (Bio_Amb and Bio_Inc) and from MES experiments: BES_Ae + Bio_Amb, BES_Ae + Bio_Inc, BES_An + Bio_Amb, and BES_An + Bio_Inc. Ambient temperatures were fluctuated between 18–25 °C, whereas incubator temperature was maintained at 30 °C. MES experiments were conducted under aerobic (Ae) and anaerobic (An) conditions.

The presence and relative enrichment of *S. ovata* in Bio_Inc provides a compelling mechanistic explanation for its superior CO_2_CE and CE. *S. ovata* is the highest-performing acetogenic biocatalyst identified to date in pure-culture MES systems, with documented acetate production rates of 51.1 g m^-2^ d^-1^ ^10^, enabled by its exceptionally high affinity for cathodic electrons and H_2_, as well as an efficient WL pathway supported by the *Sporomusa*-type Nfn transhydrogenase ^26^. In mixed-culture contexts, *S. ovata* likely functions as a keystone acetogen whose metabolic activity and electron uptake capacity disproportionately drive community-level productivity.

In aerobic BES reactors, community dynamics during days 1–5 reflected transient microaerophilic conditions (Figure 3). BES_Ae+Bio_Amb and BES_Ae+Bio_Inc showed accumulation of iso-butyrate, coinciding with the presence of oxygen-tolerant taxa, including *Bacillus*, *Pseudomonas*, and *Acinetobacter* (Figure 6)^47, 73^. Critically, *Rhodopseudomonas* was identified as a persistent and metabolically active constituent of both aerobic BES communities (17.78% in BES_Ae+Bio_Amb and 12.75% in BES_Ae+Bio_Inc, (Figure 6). This genus exhibits metabolic versatility, capable of CO_2_ fixation via the Calvin–Benson– Bassham (CBB) cycle and direct electrode-based electron uptake; thereby, partially sustaining carbon fixation under microaerophilic conditions that would otherwise suppress obligate anaerobes.^74–76^ Its persistence likely buffered against complete loss of system productivity during the oxygen-exposed early phase^77, 78^. As anoxic conditions were restored by day 5, obligate anaerobes in the community regained dominance. *Clostridium sensu* stricto 12 reached 21.87% in BES_Ae+Bio_Amb and 13.35% in BES_Ae+Bio_Inc (Figure 6), correlating with the recovery of acetate production.

Under strict anaerobiosis, BES_An+Bio_Inc exhibited approximately double the genus-level diversity of BES_An+Bio_Amb, with *Clostridium sensu* stricto 12 (27.36%) and *Rhodopseudomonas* (17.20%) as dominant taxa. BES_An+Bio_Amb was dominated by *Clostridium sensu* stricto 10 (12.1%) and *Clostridium sensu* stricto 12 (10.35%) (Figure 6). The higher diversity in BES_An+Bio_Inc likely reflects combined effects of the preservation of obligate anaerobic populations under oxygen-free conditions and optimised biofilm proliferation on electrode surfaces at 30 °C^79^. The broad product profile of BES_An+Bio_Inc, spanning seven organic acids including hexanoate and valerate (Figure 4), corroborates a functionally versatile, chain-elongating community operating at high electrochemical efficiency^72^.

Collectively, these community-level findings demonstrate that the systematic biocatalyst design strategy employed in this study, specifically the differentiation of maintenance temperature to modulate community composition, constitutes a tractable engineering lever for MES optimisation. The strong correlation between *S. ovata* enrichment, electron transfer efficiency, and acetate productivity across BES_An+Bio_Inc conditions provides mechanistic validation for targeted community design as an alternative or complement to electrode material engineering^58, 59, 80^. The anaerobic conditions for BES reactors supported a more robust product profile (Figure 4), reflecting efficient electron uptake and CO_2_ fixation. These results indicate that strictly anaerobic conditions support the activity of autotrophic and electrotrophic microbes, enhancing electron uptake from the cathode, CO_2_ fixation into acetate, and the overall BES performance while minimising side reactions.

Overall, these microbiome structures demonstrate that maintenance conditions and BES operational environments strongly shape microbial community composition and activity. Higher biomass and robust acetogen dominance resulted from maintenance at a constant 30 °C, while maintenance at ambient temperature promoted resilient, stress-adapted communities. In the context of the BES experiment, oxygen exposure transiently altered community dynamics, allowing facultative aerobes, such as *Rhodopseudomonas*, to contribute to CO_2_ fixation, whereas strictly anaerobic conditions enhanced the growth of obligate anaerobes and optimised product formation. Collectively, these findings underscore the crucial role of environmental control in MES systems in maximising microbial activity, CO_2_ conversion, and the selective production of target compounds.

## Conclusion

In this study, microbial communities of EAB were systematically enriched and maintained from activated sludge sourced from a municipal wastewater treatment facility, and their integration into BES reactors under strictly anaerobic conditions during MES conferred several operational advantages. Most notably, the low external applied potentials required demonstrated the economic viability of this approach for the reductive fixation of CO_2_ into CCCs such as organic acids, while stable and sustained production of CCCs was achieved throughout the 10-day experimental period. The findings presented here establish a clear proof of concept for systematic biocatalyst design as a strategy for improving CO_2_CE and CE in BES reactors during MES, without requiring costly electrode modification or elevated temperature operation. Collectively, these results underscore the pivotal role of optimised EAB enrichment strategies and precise BES operating conditions in driving robust and efficient MES processes. Furthermore, they highlight the considerable potential of this integrated approach for scalable industrial carbon capture and utilisation (CCU) applications, offering a promising pathway toward the biological valorisation of CO_2_ into value-added products.

## Acknowledgments

AA gratefully acknowledges the doctoral scholarship from the Indonesia Endowment Fund for Education Agency and Beasiswa Indonesia Bangkit (BIB) from the Ministry of Religious Affairs Indonesia, and MAI acknowledges support from the Environmental Biotechnology Network (EBNet) PoC grant (POC202311).

## Author contributions

A. Abbas, (Conceptualisation, experiments/investigation, and writing–original draft, writing– review, bioinformatic analysis, visualisation, and editing) and M. Ahsanul Islam (Conceptualisation, investigation, writing–review, and editing).

