## Supplementary information for "Enhancing Carbon Conversion Efficiency and Product Yield Through Systematic Biocatalyst Design for Microbial Electrosynthesis"

Leicestershire, LE11 3TU UK

**Supplementary Tables**

**Table S1. Average weekly product formation for electroactive biocatalysts maintained under different temperature conditions.**

| **Maintenance**  **conditions** | **Temperature**  **(°C)** | **Formate**  **(mg L⁻¹ week⁻¹)** | **Iso-butyrate**  **(mg L⁻¹ week⁻¹)** |
| --- | --- | --- | --- |
| Bio_Amb | 18 – 25 | 8.5 ± 1.7 | 12.5 ± 5.0 |
| Bio_Inc | 30 | 13.3 ± 4.1 | 9.1 ± 3.7 |

**Table S2. Summary of acetate production and organic acid diversity in BES_An reactors using different biocatalysts and applied potentials.**

| **Reactor conditions** | **Applied Potential (mV vs Ag/AgCl)** | **Peak Acetate concentration**  **(mg L⁻¹)** | **Acetate production rate**  **(mg L⁻¹ d⁻¹)** | **Number of organic acids detected** |
| --- | --- | --- | --- | --- |
| BES_An+Bio_Amb | –1000 | 177.03 ± 30.24 | 17.70 | 4 |
| BES_An+Bio_Amb | –600 | 90.10 ± 11.74 | 9.75 | 5 |
| BES_An+Bio_Inc | –1000 | 394.21 ± 4.01 | 39.20 | 7 |
| BES_An+Bio_Inc | –600 | 269.21 ± 4.66 | 26.42 | 4 |
| OCP+Bio_Inc | 0 (OCP) | 16.9 | – | 2 |
| OCP+Bio_Amb | 0 (OCP) | 17.1 | – | 2 |
| Control  (No biocatalysts) | –1000 | 0 | – | 0 |

**Table S3. Summary of t-test results comparing anaerobic and aerobic MES systems under varying applied potentials and biocatalysts maintenance methods.**

| **Applied Potential**  **(mV vs Ag/AgCl)** | **Biocatalyst condition** | **Comparison** | **p-value** | **Significance level** |
| --- | --- | --- | --- | --- |
| –1000 | Bio_Inc | BES_An+Bio_Inc  Vs  BES_Ae+Bio_Inc | < 0.01 | Highly significant |
| –1000 | Bio_Amb | BES_An+Bio_Amb  Vs  BES_Ae+Bio_Amb | < 0.01 | Highly significant |
| –600 | Bio_Inc | BES_An+Bio_Inc  Vs  BES_Ae+Bio_Inc | < 0.01 | Highly significant |
| –600 | Bio_Amb | BES_An+Bio_Amb  Vs  BES_Ae+Bio_Amb | < 0.01 | Highly significant |

**Table S4. Comparative analysis of the MES system performances for acetate production from CO_2_ using carbon-based cathodes.**

| **Study** | **Inoculum** | **Cathode** | **Potential (mV vs Ag/AgCl)** | **Acetate concentration (mg L^-1^)** | **Coulombic efficiency**  **(CE, %)** | **CO_2_ conversion efficiency (CO₂CE, %)** |
| --- | --- | --- | --- | --- | --- | --- |
| This study (BES_An+Bio_Inc) | Activated sludge (enriched) | Untreated carbon cloth | −1000 | 394.21 ± 4.01 | 80 | 88.11 |
| This study (BES_An+Bio_Amb) | Activated sludge (enriched) | Untreated carbon cloth | −1000 | 177.03 ± 30.24 | 51 | 31.98 |
| Tahir et al. (2020)* | Mixed culture | Untreated carbon cloth | −1000 | n. r. | 41 | n. r. |
| Zhang et al. (2013)* | Mixed culture | Untreated carbon cloth | −1000 | n. r. | 76 | n. r. |
| Jourdin et al. (2016) * | Mixed culture | NanoWeb-RVC (modified) | −585 (SHE) | n. r. | ~100 | n. r. |
| Chu et al. (2023) * | Mixed culture | Flow-electrode + PAC | n. r. | n. r. | 43.5 | n. r. |
| Bian et al. (2024)* | Mixed culture | Ni-foam carbon felt | −890 | 12,500 | n. r. | n. r. |
| Mateos et al. (2019)* | Mixed culture | Carbon felt | −800 | 1957 | 50 | 80 |

*Values from published literature; n. r. = Not reported.

**Table S5. Recorded data for Coulombic Efficiency (CE) calculation for the BES_Ae + Bio_Amb (-600 mV) configuration.**

| Day | Current1 | Current2 | Current3 | Average | Error | CCCs (molar) | M | n | F | V | %CE |
| --- | --- | --- | --- | --- | --- | --- | --- | --- | --- | --- | --- |
| 1 | 0.0041 | 0.0045 | 0.0046 | 0.0044 | 0.0003 | 0.1350 | 59.0440 | 0.0023 | 96485 | 0.1000 | 11.4637 |
| 2 | 0.0198 | 0.0205 | 0.0209 | 0.0204 | 0.0006 | 0.3177 | 59.0440 | 0.0054 | 96485 | 0.1000 | 13.6906 |
| 3 | 0.1080 | 0.1210 | 0.1250 | 0.1180 | 0.0089 | 0.8343 | 59.0440 | 0.0141 | 96485 | 0.1000 | 16.3270 |
| 4 | 0.2140 | 0.2310 | 0.2390 | 0.2280 | 0.0128 | 1.2617 | 59.0440 | 0.0214 | 96485 | 0.1000 | 19.3225 |
| 5 | 0.5230 | 0.5430 | 0.5570 | 0.5410 | 0.0171 | 2.0320 | 59.0440 | 0.0344 | 96485 | 0.1000 | 21.1231 |
| 6 | 2.5250 | 2.5750 | 2.5830 | 2.5610 | 0.0314 | 4.6683 | 59.0440 | 0.0791 | 96485 | 0.1000 | 23.5517 |
| 7 | 3.4710 | 3.6710 | 3.8410 | 3.6610 | 0.1852 | 5.5287 | 59.0440 | 0.0936 | 96485 | 0.1000 | 23.1073 |
| 8 | 4.1480 | 4.4530 | 4.6320 | 4.4110 | 0.2447 | 6.6270 | 59.0440 | 0.1122 | 96485 | 0.1000 | 27.5553 |
| 9 | 5.0970 | 5.2850 | 5.4310 | 5.2710 | 0.1674 | 7.7477 | 59.0440 | 0.1312 | 96485 | 0.1000 | 31.5179 |
| 10 | 5.3790 | 5.5650 | 5.7450 | 5.5630 | 0.1830 | 8.0850 | 59.0440 | 0.1369 | 96485 | 0.1000 | 32.5207 |

**Table S6. Recorded data for Coulombic Efficiency (CE) calculation for the BES_Ae + Bio_Amb (-1000 mV) configuration.**

| Day | Current1 | Current2 | Current3 | Average | Error | CCCs (molar) | M | n | F | V | %CE |
| --- | --- | --- | --- | --- | --- | --- | --- | --- | --- | --- | --- |
| 1 | 0.005 | 0.004 | 0.004 | 0.004 | 0.001 | 0.146 | 59.044 | 0.002 | 96485.000 | 0.100 | 13.500 |
| 2 | 0.430 | 0.290 | 0.210 | 0.310 | 0.111 | 1.301 | 59.044 | 0.022 | 96485.000 | 0.100 | 15.119 |
| 3 | 0.510 | 0.370 | 0.470 | 0.450 | 0.072 | 1.669 | 59.044 | 0.028 | 96485.000 | 0.100 | 17.139 |
| 4 | 0.435 | 0.398 | 0.431 | 0.421 | 0.020 | 1.725 | 59.044 | 0.029 | 96485.000 | 0.100 | 19.562 |
| 5 | 1.254 | 1.091 | 1.132 | 1.159 | 0.085 | 2.934 | 59.044 | 0.050 | 96485.000 | 0.100 | 20.561 |
| 6 | 1.910 | 1.760 | 1.730 | 1.800 | 0.096 | 3.742 | 59.044 | 0.063 | 96485.000 | 0.100 | 21.534 |
| 7 | 3.560 | 2.980 | 3.240 | 3.260 | 0.291 | 5.504 | 59.044 | 0.093 | 96485.000 | 0.100 | 25.719 |
| 8 | 7.210 | 6.870 | 6.410 | 6.830 | 0.401 | 8.537 | 59.044 | 0.145 | 96485.000 | 0.100 | 29.532 |
| 9 | 7.560 | 6.320 | 7.990 | 7.290 | 0.867 | 9.670 | 59.044 | 0.164 | 96485.000 | 0.100 | 35.498 |
| 10 | 11.510 | 9.680 | 9.380 | 10.190 | 1.153 | 11.102 | 59.044 | 0.188 | 96485.000 | 0.100 | 33.478 |

**Table S7. Recorded data for Coulombic Efficiency (CE) calculation for the BES_Ae + Bio_Inc (-600 mV) configuration.**

| Day | Current1 | Current2 | Current3 | Average | Error | CCCs (molar) | M | n | F | V | %CE |
| --- | --- | --- | --- | --- | --- | --- | --- | --- | --- | --- | --- |
| 1 | 0.002 | 0.003 | 0.002 | 0.002 | 0.000 | 0.113 | 59.044 | 0.002 | 96485.000 | 0.100 | 15.365 |
| 2 | 0.007 | 0.008 | 0.008 | 0.008 | 0.001 | 0.243 | 59.044 | 0.004 | 96485.000 | 0.100 | 21.444 |
| 3 | 0.048 | 0.043 | 0.086 | 0.059 | 0.024 | 0.741 | 59.044 | 0.013 | 96485.000 | 0.100 | 25.757 |
| 4 | 0.137 | 0.142 | 0.168 | 0.149 | 0.017 | 1.174 | 59.044 | 0.020 | 96485.000 | 0.100 | 25.601 |
| 5 | 0.421 | 0.378 | 0.386 | 0.395 | 0.023 | 2.055 | 59.044 | 0.035 | 96485.000 | 0.100 | 29.589 |
| 6 | 1.652 | 1.487 | 1.454 | 1.531 | 0.106 | 4.422 | 59.044 | 0.075 | 96485.000 | 0.100 | 35.343 |
| 7 | 2.753 | 2.438 | 2.738 | 2.643 | 0.178 | 5.471 | 59.044 | 0.093 | 96485.000 | 0.100 | 31.347 |
| 8 | 2.951 | 2.765 | 2.852 | 2.856 | 0.093 | 6.135 | 59.044 | 0.104 | 96485.000 | 0.100 | 36.474 |
| 9 | 4.152 | 4.045 | 4.196 | 4.131 | 0.078 | 7.676 | 59.044 | 0.130 | 96485.000 | 0.100 | 39.472 |
| 10 | 4.761 | 4.038 | 3.654 | 4.151 | 0.562 | 8.070 | 59.044 | 0.137 | 96485.000 | 0.100 | 43.425 |

**Table S8. Recorded data for Coulombic Efficiency (CE) calculation for the BES_Ae + Bio_Inc (-1000 mV) configuration.**

| Day | Current1 | Current2 | Current3 | Average | Error | CCCs (molar) | M | n | F | V | %CE |
| --- | --- | --- | --- | --- | --- | --- | --- | --- | --- | --- | --- |
| 1 | 0.003 | 0.003 | 0.003 | 0.003 | 0.000 | 0.125 | 59.044 | 0.002 | 96485.000 | 0.100 | 15.444 |
| 2 | 0.257 | 0.246 | 0.247 | 0.250 | 0.006 | 1.344 | 59.044 | 0.023 | 96485.000 | 0.100 | 20.007 |
| 3 | 0.795 | 0.987 | 0.877 | 0.890 | 0.096 | 2.754 | 59.044 | 0.047 | 96485.000 | 0.100 | 23.580 |
| 4 | 1.368 | 1.721 | 1.507 | 1.530 | 0.178 | 3.904 | 59.044 | 0.066 | 96485.000 | 0.100 | 27.570 |
| 5 | 2.317 | 2.153 | 2.415 | 2.290 | 0.132 | 5.000 | 59.044 | 0.085 | 96485.000 | 0.100 | 30.218 |
| 6 | 4.728 | 3.913 | 4.947 | 4.530 | 0.545 | 6.963 | 59.044 | 0.118 | 96485.000 | 0.100 | 29.624 |
| 7 | 6.312 | 6.613 | 5.723 | 6.220 | 0.453 | 8.619 | 59.044 | 0.146 | 96485.000 | 0.100 | 33.052 |
| 8 | 6.861 | 6.432 | 6.905 | 6.730 | 0.261 | 9.525 | 59.044 | 0.161 | 96485.000 | 0.100 | 37.310 |
| 9 | 8.157 | 8.117 | 8.428 | 8.235 | 0.169 | 10.920 | 59.044 | 0.185 | 96485.000 | 0.100 | 40.077 |
| 10 | 9.865 | 9.856 | 9.654 | 9.876 | 0.119 | 11.976 | 59.044 | 0.203 | 96485.000 | 0.100 | 40.193 |

**Table S9. Recorded data for Coulombic Efficiency (CE) calculation for the BES_An + Bio_Amb (-600 mV) configuration.**

| Day | Current1 | Current2 | Current3 | Average | Error | CCCs (molar) | M | n | F | V | %CE |
| --- | --- | --- | --- | --- | --- | --- | --- | --- | --- | --- | --- |
| 1 | 0.003 | 0.003 | 0.003 | 0.003 | 0.000 | 0.147 | 59.044 | 0.002 | 96485.000 | 0.100 | 21.513 |
| 2 | 2.765 | 2.987 | 3.149 | 2.967 | 0.193 | 4.693 | 59.044 | 0.079 | 96485.000 | 0.100 | 20.547 |
| 3 | 43.560 | 46.310 | 45.460 | 45.110 | 1.408 | 20.396 | 59.044 | 0.345 | 96485.000 | 0.100 | 25.523 |
| 4 | 22.580 | 25.350 | 24.410 | 24.110 | 1.409 | 15.477 | 59.044 | 0.262 | 96485.000 | 0.100 | 27.497 |
| 5 | 186.530 | 200.050 | 210.120 | 198.900 | 11.837 | 46.842 | 59.044 | 0.793 | 96485.000 | 0.100 | 30.532 |
| 6 | 16.450 | 20.030 | 17.550 | 18.010 | 1.834 | 15.212 | 59.044 | 0.258 | 96485.000 | 0.100 | 35.560 |
| 7 | 1.125 | 1.542 | 1.482 | 1.383 | 0.225 | 4.478 | 59.044 | 0.076 | 96485.000 | 0.100 | 40.135 |
| 8 | 12.645 | 15.675 | 16.237 | 14.859 | 1.932 | 15.282 | 59.044 | 0.259 | 96485.000 | 0.100 | 43.497 |
| 9 | 0.012 | 0.014 | 0.012 | 0.012 | 0.001 | 0.452 | 59.044 | 0.008 | 96485.000 | 0.100 | 45.668 |
| 10 | 1.146 | 1.268 | 1.171 | 1.195 | 0.064 | 4.621 | 59.044 | 0.078 | 96485.000 | 0.100 | 49.463 |

**Table S10. Recorded data for Coulombic Efficiency (CE) calculation for the BES_An + Bio_Amb (-1000 mV) configuration.**

| Day | Current1 | Current2 | Current3 | Average | Error | CCCs (molar) | M | n | F | V | %CE |
| --- | --- | --- | --- | --- | --- | --- | --- | --- | --- | --- | --- |
| 1 | 0.003 | 0.002 | 0.004 | 0.003 | 0.001 | 0.154 | 59.044 | 0.003 | 96485.000 | 0.100 | 23.526 |
| 2 | 4.890 | 5.010 | 4.230 | 4.710 | 0.420 | 6.896 | 59.044 | 0.117 | 96485.000 | 0.100 | 27.944 |
| 3 | 51.240 | 47.140 | 49.490 | 49.290 | 2.057 | 21.316 | 59.044 | 0.361 | 96485.000 | 0.100 | 25.513 |
| 4 | 98.351 | 95.036 | 97.883 | 96.643 | 1.794 | 35.046 | 59.044 | 0.594 | 96485.000 | 0.100 | 35.173 |
| 5 | 165.230 | 147.230 | 166.070 | 169.515 | 10.643 | 43.751 | 59.044 | 0.741 | 96485.000 | 0.100 | 31.252 |
| 6 | 48.571 | 45.654 | 48.371 | 46.649 | 1.629 | 27.032 | 59.044 | 0.458 | 96485.000 | 0.100 | 43.354 |
| 7 | 21.321 | 19.268 | 19.849 | 20.146 | 1.058 | 17.620 | 59.044 | 0.298 | 96485.000 | 0.100 | 42.653 |
| 8 | 4.761 | 4.143 | 4.791 | 4.569 | 0.366 | 8.841 | 59.044 | 0.150 | 96485.000 | 0.100 | 47.347 |
| 9 | 10.258 | 11.287 | 6.580 | 9.211 | 2.475 | 12.272 | 59.044 | 0.208 | 96485.000 | 0.100 | 45.251 |
| 10 | 5.476 | 4.358 | 9.971 | 6.951 | 2.971 | 11.288 | 59.044 | 0.191 | 96485.000 | 0.100 | 50.734 |

**Table S11. Recorded data for Coulombic Efficiency (CE) calculation for the BES_An + Bio_Inc (-600 mV) configuration.**

| Day | Current1 | Current2 | Current3 | Average | Error | CCCs (molar) | M | n | F | V | %CE |
| --- | --- | --- | --- | --- | --- | --- | --- | --- | --- | --- | --- |
| 1 | 0.231 | 0.211 | 0.284 | 0.242 | 0.038 | 1.767 | 59.044 | 0.030 | 96485.000 | 0.100 | 35.721 |
| 2 | 0.870 | 0.760 | 1.250 | 0.960 | 0.257 | 3.604 | 59.044 | 0.061 | 96485.000 | 0.100 | 37.446 |
| 3 | 17.432 | 15.434 | 17.237 | 16.713 | 1.102 | 16.630 | 59.044 | 0.282 | 96485.000 | 0.100 | 45.797 |
| 4 | 66.325 | 67.231 | 61.798 | 65.118 | 2.911 | 33.996 | 59.044 | 0.576 | 96485.000 | 0.100 | 49.121 |
| 5 | 31.415 | 30.043 | 31.167 | 30.875 | 0.731 | 25.233 | 59.044 | 0.427 | 96485.000 | 0.100 | 57.074 |
| 6 | 32.657 | 31.467 | 28.495 | 30.873 | 2.144 | 26.710 | 59.044 | 0.452 | 96485.000 | 0.100 | 63.954 |
| 7 | 1.941 | 1.897 | 1.793 | 1.877 | 0.076 | 6.772 | 59.044 | 0.115 | 96485.000 | 0.100 | 67.614 |
| 8 | 13.456 | 12.435 | 8.747 | 11.546 | 2.477 | 16.987 | 59.044 | 0.288 | 96485.000 | 0.100 | 69.171 |
| 9 | 53.378 | 52.145 | 62.438 | 55.987 | 5.621 | 39.119 | 59.044 | 0.663 | 96485.000 | 0.100 | 75.648 |
| 10 | 4.214 | 4.213 | 3.561 | 3.996 | 0.377 | 10.300 | 59.044 | 0.174 | 96485.000 | 0.100 | 73.478 |

**Table S12. Recorded data for Coulombic Efficiency (CE) calculation for the BES_An + Bio_Inc (-1000 mV) configuration.**

| Day | Current1 | Current2 | Current3 | Average | Error | CCCs (molar) | M | n | F | V | %CE |
| --- | --- | --- | --- | --- | --- | --- | --- | --- | --- | --- | --- |
| 1 | 0.331 | 0.515 | 0.447 | 0.431 | 0.093038 | 2.18233 | 59.044 | 0.036961 | 96485 | 0.1 | 30.58242 |
| 2 | 41.178 | 37.651 | 42.154 | 40.328 | 2.368872 | 22.80433 | 59.044 | 0.386226 | 96485 | 0.1 | 35.68915 |
| 3 | 25.675 | 21.342 | 27.865 | 24.973 | 3.319652 | 19.86034 | 59.044 | 0.336365 | 96485 | 0.1 | 43.71303 |
| 4 | 88.943 | 74.543 | 89.463 | 84.286 | 8.467947 | 35.59166 | 59.044 | 0.602799 | 96485 | 0.1 | 41.59578 |
| 5 | 24.357 | 17.354 | 25.342 | 22.357 | 4.355464 | 20.837 | 59.044 | 0.352906 | 96485 | 0.1 | 53.74835 |
| 6 | 51.389 | 41.355 | 49.978 | 47.571 | 5.431823 | 32.52667 | 59.044 | 0.550889 | 96485 | 0.1 | 61.55243 |
| 7 | 107.385 | 87.475 | 113.476 | 102.779 | 13.59878 | 49.25833 | 59.044 | 0.834265 | 96485 | 0.1 | 65.33761 |
| 8 | 512.668 | 435.775 | 529.254 | 492.576 | 49.87643 | 115.112 | 59.044 | 1.949597 | 96485 | 0.1 | 74.45197 |
| 9 | 40.156 | 31.267 | 38.315 | 36.577 | 4.6918 | 31.99734 | 59.044 | 0.541924 | 96485 | 0.1 | 77.469 |
| 10 | 4.156 | 2.252 | 5.397 | 3.945 | 1.584104 | 10.44833 | 59.044 | 0.176958 | 96485 | 0.1 | 76.58699 |

Note:	Current	=	Ampere (A)

CCCs	=	Commodity chemicals concentration (M)

M	=	Molar mass

n	=	Number of moles of electrons recovered into CCCs

F	=	Faraday constant (C/mol)

V	=	reactor volume (L)

**Supplementary Figure**

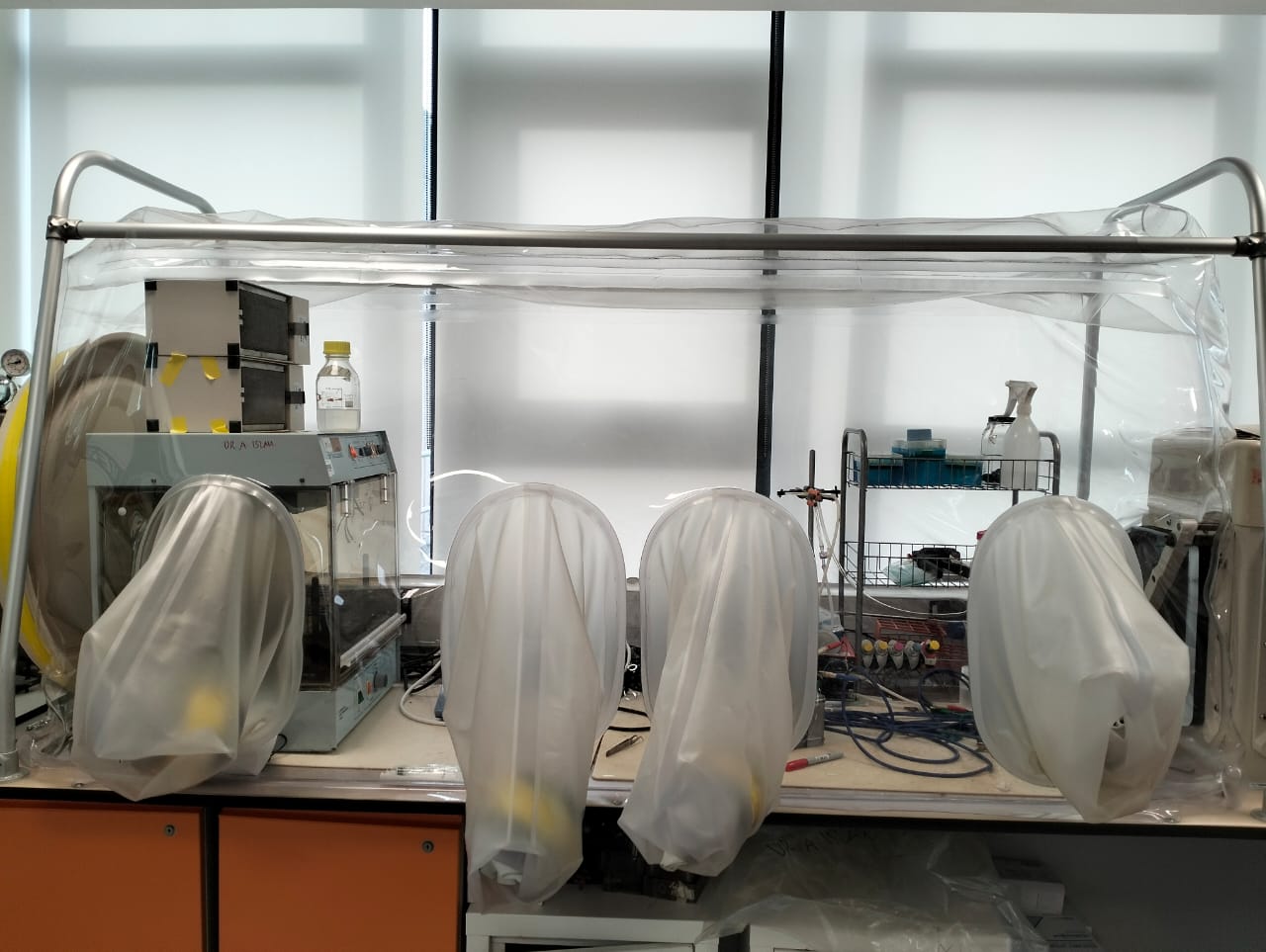

Figure S1. Vinyl anaerobic chamber for maintaining strictly anaerobic conditions during BES operation and sample handling. The system includes glove ports, controlled gas atmosphere, and an enclosed workspace for anaerobic manipulation.
